# Long non-coding RNA *hsr-omega* provides scaffolding for the nuclear domain B-body

**DOI:** 10.1101/2025.05.07.652789

**Authors:** SooBin An, Sharar Haque, Miranda Adams, Luis Enrique Torres Rodriguez, Kaveh Kiani, Anton L. Bryantsev

## Abstract

Nuclear domains (NDs)—such as nucleoli or nuclear speckles—are membraneless, organelle-like compartments that concentrate and retain nuclear proteins. Despite their ubiquitous presence in the cell, the organization, regulation, and functions of many NDs remain poorly understood. The B-body is a prominent nuclear domain observed in developing flight muscles of *Drosophila*. In this study, we expand the understanding of B-body composition and function. We identify several additional RNA-binding proteins (RBPs) as B-body components and show that some proteins can dynamically disappear from this ND. We further demonstrate that the B-body contains an RNA component, which was identified as the long non-coding RNA *hsrω*. Genetic analyses reveal that *hsrω* acts as a structural scaffold for the B-body, and its depletion leads to B-body disassembly. In contrast, loss of the resident protein Bruno (Bru), a splicing factor, does not compromise B-body integrity. Finally, we show that imbalance in the *hsrω*/Bru ratio promotes Bru aggregation, suggesting that the B-body plays a role in maintaining nuclear protein homeostasis.

## Introduction

Nuclear domains (NDs) are organelle-like structures within the cell nucleus which do not have membranes yet can selectively accumulate and retain certain nuclear factors. Depending on their composition, NDs are different in abundance, distribution, shape, and size. The nucleolus is the most prominent and recognizable among NDs; other examples include Cajal bodies, nuclear speckles, PML bodies, *etc*. [1]. Although some NDs have been known for a long time (*e*.*g*., Cajal bodies [2]), their formation and regulation remain elusive.

B-body is a novel nuclear domain that was described in developing flight muscles in *Drosophila* [3]. Indirect flight muscles (IFM), consisting of Dorsoventral (DVM) and dorsal longitudinal (DLM) muscles, are highly specialized muscles in adult flies that belong to the fibrillar muscle type and are strikingly different from the rest of adult muscles [4]. During metamorphosis, DLMs are formed on the basis of pre-existing Larval Oblique Muscles (LOMs) that serve as a fusion template for numerous myoblasts from 16 h to 24 h after puparium formation (apf). During that developmental time frame, B-bodies are the most prominently visible in LOM’s nuclei [3].

B-bodies contain Bruno (Bru, encoded by the gene *bru1* (FBgn0000114)) and Muscleblind (Mbl), which both are RNA-binding proteins (RBPs) that regulate pre-mRNA splicing [3]. In particular, Bru enables IFM-specific isoforms of muscle structural genes and is critical for the unique appearance of flight muscles [3]. Besides flight muscles, Bru is expressed in gonads, where it mediates mRNA transport and translational silencing [5]. Bru binds Bruno-response elements (BREs) at the 3’UTRs of specific ovarian transcripts (e.g., *osk, gurk*, etc) to prevent their premature translation in the developing oocyte. The sequence preference for Bru binding was established based on BRE and *in vitro* binding assays [5].

The principal component of the B-body, protein Bruno, is capable of diverse intermolecular interactions. Bru can self-dimerize and bind other proteins directly; additional proteins can cooperate with Bru via binding to the common RNA targets [6]. Such oligomerizing activity enables Bru to assemble into large complexes, “silencing particles”, that contain mRNA oligomers and other proteins [7]. Concomitantly, Bru is present in the perinuclear organelle-like structures “nuage” that can be detected by microscopy in germline cells [8]. Whether Bru can serve as a critical structural element for B-bodies or other structures where it has been detected is not clear.

Long non-coding RNA (lncRNA) is a diverse group of transcripts that can play a range of functions in the cell [9]. lncRNA as a structural component is present in several NDs. For example, lncRNA Xist is directly involved in the formation of the Barr body, which represents an inactivated and structurally condensed extra copy of the X-chromosome in females [10]. Similarly, lncRNA NEAT1 is a critical structural component of paraspeckles, a set of NDs hosting RBPs involved in RNA editing and splicing, which are located adjacent to classical nuclear speckles [11]. In *Drosophila*, the lncRNA *hsrω* forms distinct nuclear speckles that accumulate several types of hnRNPs [12]. However, not all NDs have an RNA scaffold. To assemble PML bodies, protein modification by SUMOylation is sufficient [13]. Although many NDs, such as Cajal bodies, histone locus body, or the nucleolus are packed with short- and medium-size RNA that is part of ribonucleoprotein complexes concentrating at those locations, there is no evidence of lncRNA presence [14]. Similarly, no structural lncRNA exists in nuclear speckles as these NDs are resistant to RNase treatment [15]. Therefore, the role of lncRNA as a scaffold in NDs is not universal and how it is selected to be involved in some structures but not the other is of if there are any rules about why it is required for some NDs but not the other is currently unknown.

The *Drosophila* ncRNA gene *heat stress response omega (hsrω)* is a Drosophila lncRNA that was discovered as a puff on a polytene chromosome in the cytoband 93D that strongly activated by heat shock and has a long history of research [16]. The hsrω locus produces multiple transcripts that can be categorized into three groups: small and often spliced transcripts under 3 kb in length (isoforms RA, RC, RD, RH, by Flybase annotations), longer 15-kb transcripts RB and RG, and a single, long, non-spliced isoform RF, extending beyond 21 kb. The longer isoforms have a unique repetitive region spanning more than 10 kb and consisting of an array of direct repeats [10]. From early studies, it was evident that *hsrω* is non-coding and associated with RNPs [16]. In the interphase nucleus, hsr forms highly dynamic NDs known as omega speckles. These structures contain various RBPs and can be found in many types of cells and tissues [16]. Hsr has been implicated in stress protection by sheltering splicing factors upon heat shock [17], but it’s function during development is poorly understood.

In this study, we analyzed molecular composition and organization of the B-body. We have expanded the list of B-body proteins besides Bru to show that some proteins accumulate there transiently or partially. We have identified lncRNA *hsrω* within B-bodies and ran isoform analysis to show the longest RF isoform as B-body resident. We further reveal that *hsrω* is the essential scaffolding component of the B-body. Finally, we demonstrate that reduction of *hsrω* leads to spontaneous protein aggregation of Bru, suggesting the role for B-bodies as a safe depot for handling excessive amount of RPBs prior to the active phase of myogenesis.

## MATERIALS & METHODS

### Fly stocks and fly husbandry

Genetic lines for this study were obtained from the Bloomington Drosophila Stock Center (BDSC) (**Table 1**). Flies were kept on Jazz-Mix food (Fisherbrand™). Crosses and fly development were set at 25 °C.

**Table 1.** List of Drosophila genetic lines used in this study. BDSC – Bloomington Drosophila Stock Center, KDSC – Kyoto Drosophila Stock Center.

| ID | Line Name | Genotype | Source |
| --- | --- | --- | --- |
| 3605 | w[1118] | w[1118] | BDSC |
| 50742 | Mef2-Gal4 | w[*]; P{w[+m*]=Mef2-GAL4.247}3 | BDSC |
| 36034 | attP40 | y[1] w[*];<br>Mi{y[+mDint2]=MIC}ple[MI02717]/TM<br>3, Sb[1] Ser[1] | BDSC |
| 44483 | Bru-IR | y[1] v[1]; P{y[+t7.7]<br>v[+t1.8]=TRiP.HMC02374}attP2 | BDSC |
| Sandy | GFP::Bru(WT) | w*; P{w[+mW UAS-GFP::Bru]} ZH-51C | (Oas *et al.*, 2014) |
| 5797 | Df(3R)GC14 | Df(3R)GC14/TM6B, Tb[+] | BDSC |
| 59617 | *hsr-ω<sup>66</sup>* | w[1118]; lncRNA:Hsromega[<sup>66</sup>]/TM6B,<br>Tb[1] | BDSC |
| 9480* | Df(3R)ED6052 | w[1118]; Df(3R)ED6052,<br>P{w[+mW.Scer<sup>FRT</sup>.hs3]=3'.RS5+3.3'}<br>ED6052/TM6B, Tb[1] | *) Modified BDSC's<br>9480 by replacing TM6C<br>with TM6B balancer. |

### Selecting lncRNA candidates for screening

Sequences of all annotated lncRNA transcripts were downloaded from FlyBase (genome assembly r6.29). The top ten short tetranucleotides preferred by Bru RNA binding motifs in vitro [18]. The selected tetranucleotides were searched in lncRNA transcripts using the regular expression (regex) tool available from the Galaxy platform (usegalaxy.org). The transcripts that were enriched (at least two copies in a 100-bp window) by each of the tetranucleotides were ranked by size. The longest transcripts were selected as top candidates for FISH screening.

### Genetic knockout of hsromega

Two different combinations of deficiencies were used. *Df(3R)ED6052*/*hsr-ω*^*66*^ and *Df(3R)ED6052*/ *Df(3R)GC14*. The *ED6052* deficiency spans 66 kb and removes the entire sequence of *hsrω. hsr-ω*^*66*^ has an excision of 1,598 bp, removing the upstream regulatory sequences of *hsrω*, including the promoter and transcription start site [19]. The *GC14* deficiency has no molecularly defined breakpoints available but experimentally was shown to render the *hsrω* gene non-functional [20]. All fly stock with these deletions were balanced against a *TM6B* balancer chromosome. Progeny resulting from crossing flies with deletions contained a mixture of genotypes. Control flies having a balancer chromosome (*TM6B*^*+*^, short pupa) and heterozygous deletion pupae (e.g., *ED6052*/*hsr-ω*^*66*^, normal long pupa) were distinguishable by their *tubby* phenotype. Based on these morphologic differences control and experimental (double deletion) white pupae were identified and transferred into a new separate vial and let develop at 25 ^°^C for a designated time (e.g., 16 h, 24 h, or 48 h) before being frozen in liquid nitrogen and analyzed.

### Sample collection and cryosectioning

Pupae, from 2 to 5, were placed on a spatula, covered with Tissue-Tek (Sakura®), and flash-frozen by submerging into liquid nitrogen. The frozen blocks were kept at -80 °C until cryosectioning. Frozen samples were moved into a styrofoam box with liquid nitrogen and brought into the cryotome machine (Epredia HM525 NX). Blocks were sectioned at a thickness of 10 μm at -20 °C from the topmost dorsal side to the bottommost ventral side. All sectioned samples were collected on a clean microscope slide. Slides were kept in a slide box at +4 °C until the staining process.

### Antibodies and dyes

Polyclonal primary antibodies against the following proteins were used: Bru1 (gift from Dr. Paul MacDonald, rabbit), Mbl (gift from Dr. Darren Moncton, sheep), and GFP (Aves Labs, chicken) (Houseley *et al*., 2005; Kanke *et al*., 2015). Monoclonal primary antibodies (Glo, Hfp, Orb2, Fl(2)d, and Rump) were purchased from Developmental Studies Hybridoma Bank (DSHB, mouse). Phalloidin conjugated with iFluor 488 was from Abcam (ab176753), and DNA dye DAPI was from Sigma-Aldrich. All custom-made RNA FISH probes were from LGC Biosearch Technologies. The final concentration of the RNA probes was 12.5 µM. *hsrω_RB, RG, RF* FISH probes were labeled with Cy5 dye, and *hsrω_RF* FISH probe contained Quasar® 670 (Q670) dye. Besides these two probes, pan-specific *hsrω* FISH probe contained Quasar® 570 (Q570) dye. The T30 FISH probe was used as a positive control to validate the staining process as it binds to the poly(A) tail of mRNAs.

**Table 2** summarizes the reagents used in this study for staining. Stock solutions were stored in small aliquots either at -20 °C (antibodies) or -80 °C (RNA probes) and were defrosted immediately before use. Stellaris® RNA FISH probes with fluorescent labels were custom designed via Stellaris Probe Designer (LGC Biosearch Technologies), based on the sequences of candidate genes downloaded from Flybase. In the situations when multiple annotated isoforms existed, the probes were designed against the regions common to all isoforms, except the situations when isoform-specific staining was desired (*hsrω* isoforms, described above).

**Table 2.**
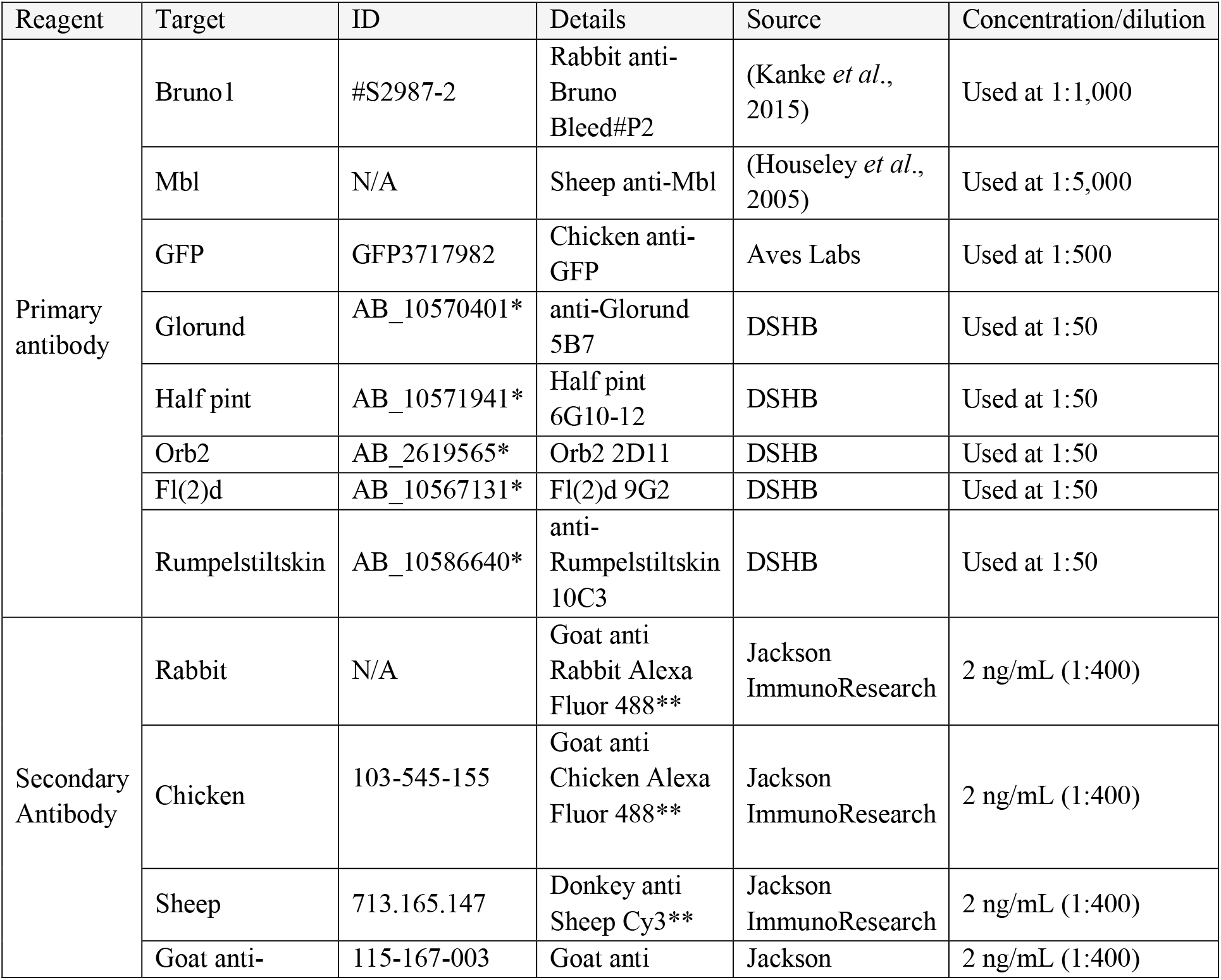
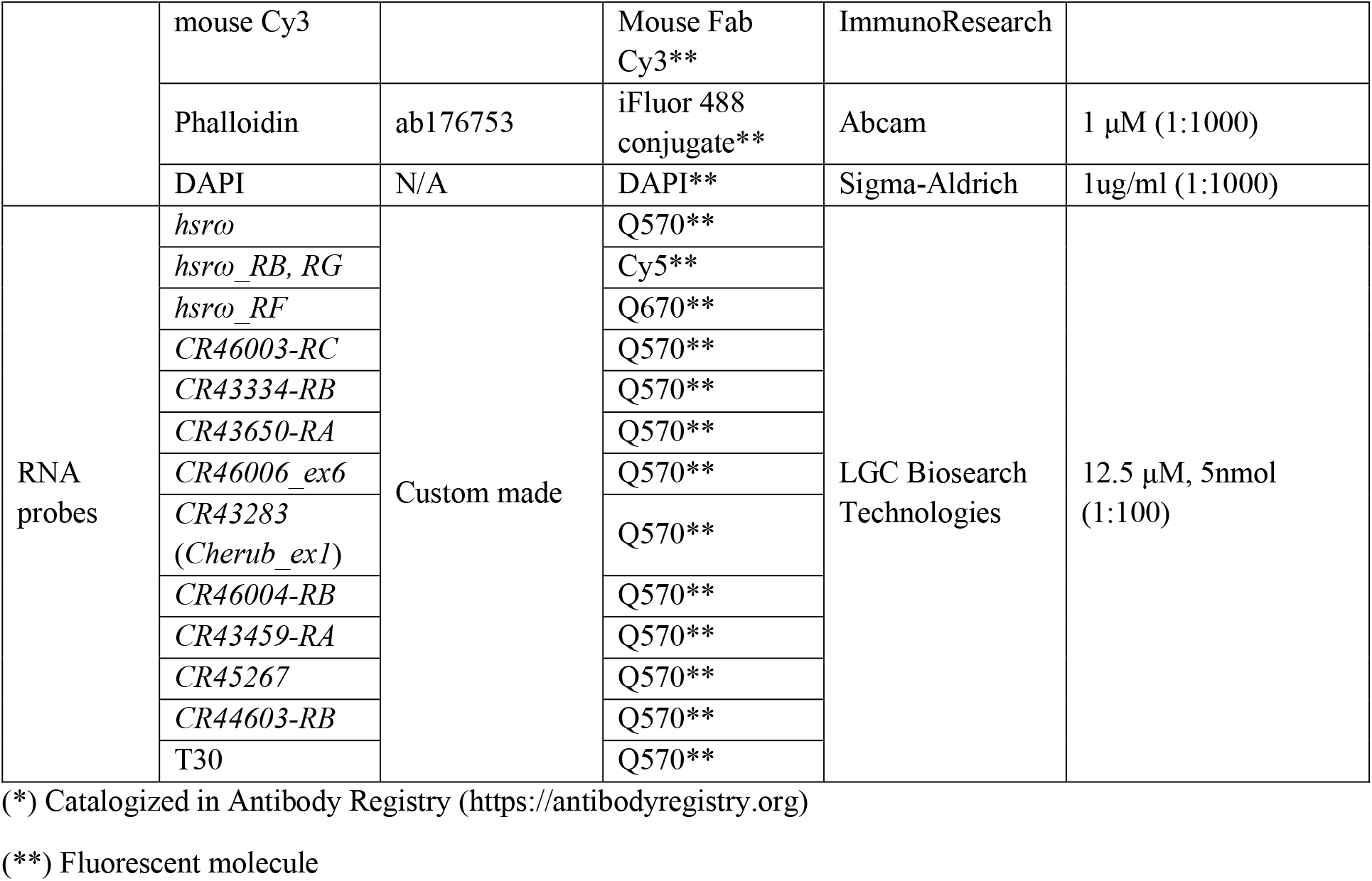
The list of staining reagents used in this study.

### Immunofluorescence staining

Slides with cryosections were fixed for 10 minutes with 3.7% formaldehyde solution prepared in phosphate buffer saline (PBS), washed three times with phosphate buffer saline containing 0.1% Triton X-100 (PBTx), and submerged into the permeabilization solution (1% Triton X-100 in PBS) for 30 minutes on a rocking platform at 90 rpm. A droplet (95 μL) of the primary staining solution (containing primary antibodies diluted in PBTx containing 1% bovine serum albumin (BSA)) was applied per slide, covered with a coverslip, placed into a humid chamber, and incubated overnight at room temperature, in the dark. On the following day, the slides were washed two times with PBTx, and incubated with 95 μL/slide of the secondary staining solution (containing secondary antibodies) with a coverslip in the humid chamber at room temperature for 1 hour in the dark. After a brief wash with PBTx, the slides were permanently mounted with a coverslip using the Mowiol medium (0.12 g/ml Mowiol 4-88 (Sigma), 0.3 g/ml glycerol, 120 mM Tris pH 8). After air-drying for 1 h, the slides were stored at 4 °C until imaging.

### Fluorescence in situ hybridization (FISH)

In order to protect RNA from degradation, all reagents that were used were of RNase-free grade; all work was conducted in a laminate flow workstation (Nuaire) Slides with cryosections were fixed with 3.7% formaldehyde, washed with PBS three times, and submerged into 70% ethanol at room temperature for an hour. After that, the slides were incubated with 500 μL Stellaris® RNA FISH Wash Buffer A for 5 minutes. Staining was done with a FISH staining solution consisting of 180 μL Stellaris® RNA FISH hybridization buffer, 20 μL formamide (Thermo Fisher Scientific), and 2 μL of each Stellaris® FISH probe (final concentration 12.5 μM) (**Table 2**). 200 μL of the probe solution was applied per slide and covered by a coverslip. The slides were incubated in the humid chamber at 37 °C overnight in the dark. On the following day, the slide was washed with Buffer A and kept in Buffer A in the humid chamber at 37 °C for 30 minutes in the dark. For nuclear counterstaining, Buffer A with 1 uM DAPI was applied and incubated at 37 °C for 30 minutes in the dark with a coverslip. Finally, the slide was rinsed with Stellaris® RNA FISH Wash Buffer B for 5 minutes after removing the coverslip. 90 μL of mounting media was applied to the slide to mount the coverslip. The mounted slides were air-dried and kept in a 4 °C slide folder before being visualized under the microscope.

### Immuno-FISH staining

We followed guidance from Kochan *et al*. (2015) for double staining, combining immunofluorescence and FISH. The entire procedure took three days, combining the protocols of immunofluorescence (days 1-2) and FISH (days 2-3) as described above, but with a few important modifications. First, BSA from the primary staining solution was replaced by PBTx, as BSA may be a source of contaminating RNases (Kochan *et al*., 2015). Second, following the last wash of the immunofluorescence protocol, the samples were post-fixed by incubating for 10 min in 3.7% formaldehyde solution. This was done to retain the antibodies during the stringent hybridization conditions.

### RNase treatment of cryosections

Cryosections from a frozen block of 1-day-old young adult flies (*88F>Act88F* KD, 25 °C) were distributed equally to three slides, which were treated differently. One slide was exempted from any treatment before the fixation. Cell lysis was performed on fresh, unfixed tissue sections for the other two slides using either cytoskeleton (CSK) buffer (10 mM Tris pH 6.8, 100 mM NaCl, 300 mM sucrose, 3 mM MgCl_2_, 1 mM EGTA, 0.5% Triton X-100) alone or in combination with RNase A (1 ug/ml) for 10 min at room temperature before fixation. After the treatment, the standard immunofluorescence staining was performed for Bru, as described above. A single-pass confocal optical section was used to collect images of 160 to 250 nuclei per slide in total. The total number of nuclei observed and the number of nuclei possessing B-body were recorded from the images by manual count.

### Microscopy

We used the Lecia MZ75 microscope for fly sorting, making frozen blocks, and counting the number of pupae. Images were taken using Zeiss Axio Imager M2 and Zeiss LSM 900 microscopes. Zeiss Axio Imager M2 is a fluorescent microscope to visualize IF/FISH double staining cells and to produce images for B-body area quantification. Zeiss LSM 900 is a confocal microscope to produce images of B-body in higher resolution. The objective used in both microscopes was the Zeiss Plan-Apochromat 63x/1.4 Oil DIC. All the camera settings, including exposure time, were fixed when investigating one slide. The z-stack function with a 0.5 µm interval was used in both microscopes to produce images of different focal planes. Final images were always produced using the maximum-intensity projection algorithm with the Zen 3.0 software.

### B-body quantification

Cryosections of control and experimental pupae were placed on the same slide to ensure equal treatment during immunostaining. Each microscope slide contained 2 to 5 pupae. B-body images were taken using Zeiss Axio Imager M2 with the 63x/1.4 Oil DIC objective. During the imaging of optical stacks, the camera settings (including the exposure time) were kept unchanged between images of the same slide. Special care was applied to keep signal intensity to 25-30% of the maximal dynamic range of the camera. 5 to 10 optical stacks were flattened in the Zen software via the Maxum Projections algorithm.

Eleven LOM nuclei from the images obtained from 2-5 control and experimental pupae were selected, respectively. An investigator assigned a random code to each cropped image, and the other investigator analyzed the images in a blind manner. B-body analyses consisted of measurement of B-body area and group identification (control and experimental). To measure the area of B-body, each image in default brightness and contrast levels was transferred to Adobe Photoshop 2023. The intensity of the image pixels was adjusted in a new layer by selecting Levels in the Adjustments menu. Then the image was converted into black-and-white images via the Threshold option, and the pixel level was adjusted to selectively show the boundary of B-bodies. B-bodies were selected by the magic wand tool in Photoshop, and the area was measured in micrometers (using a conversion scale). If a nucleus had multiple B-bodies, their areas were summed up. After identifying the control and experimental groups of B-body, the average and standard deviation of total data were calculated and visualized as a bar graph in Microsoft Excel.

### Statistical analysis

A paired Student’s *t-*test was used to assess differences of the isoform staining intensities between B-bodies and omega speckles (**Fig. 5**). An unpaired Student’s *t-*test was used to evaluate differences in B-body areas in knockout and knockdown experiments (**Figs. 6&7**). Fisher’s exact test was applied to compare the frequency of nuclear aggregates (**Fig. 8**).

## RESULTS

### B-body is a nuclear domain with dynamic protein composition

B-bodies are prominent at the onset of flight muscle development (**Fig. 1A**). At 16 h apf, the LOM templates with large polyploid nuclei are surrounded by fusion-competent myoblasts with small diploid nuclei. Shortly after, myoblasts start fusing with LOMs to form nascent myofibers of DLMs. By 24 h apf, the myoblast fusion is mainly complete, but the original LOM nuclei still can be traced within DLMs via their notably bigger size up to 48 h apf, after which all DLM nuclei become polyploid and uniform (**Fig. 1A**).

**Figure 1:**
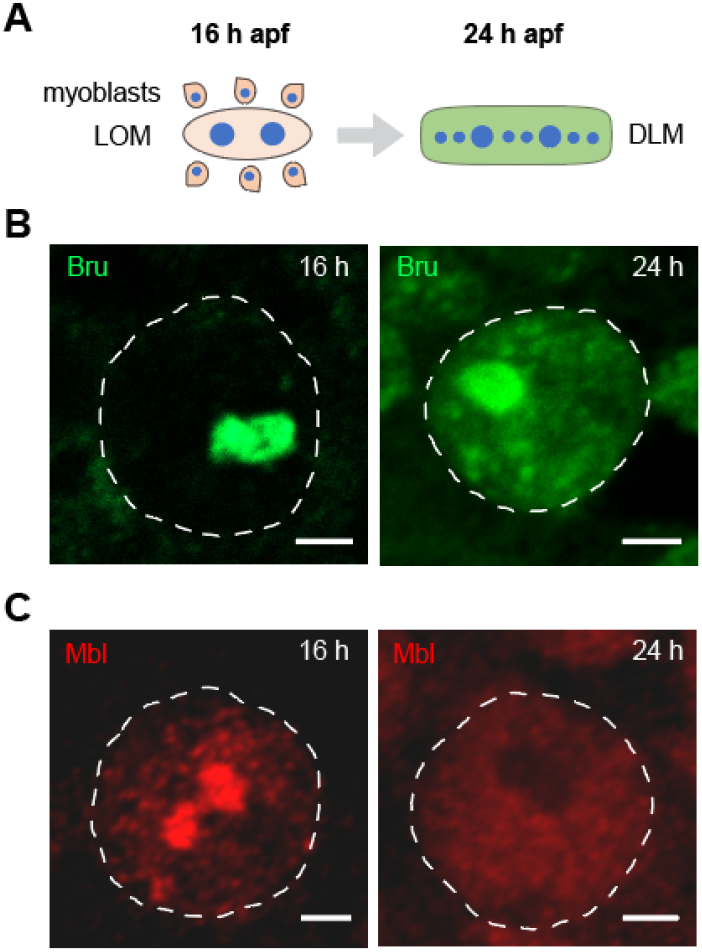
Location and appearance of B-bodies. **A**. Schematic representation of early stages of Dorsal Longitudinal Muscle (DLM) development. At 16 h after puparium formation (apf), Larval Oblique Muscle (LOM) cells with large polyploid nuclei are surrounded by fusion-competent myoblasts containing small diploid nuclei. By 24 h apf, myoblasts have fused with the LOM template to form nascent DLM fibers. LOM nuclei remain identifiable by their significantly larger size. **B**. Distribution of Bru in LOM nuclei at 16 and 24 h apf. By 24 h, Bru becomes more diffuse and forms small nuclear speckles, while the B-body remains largely unchanged. **C**. Dynamics of Mbl protein localization in B-bodies. Mbl is entirely lost from B-bodies by 24 h apf. Bru and Mbl proteins were detected by immunofluorescence using a polyclonal antibody. Dashed lines mark nuclear boundaries. Scale bars = 2 µm (all panels).

Bru localization in B-bodies is dynamic: it was present exclusively in the B-body at 16 h apf, but at 24 h apf part of Bru becomes more generally distributed throughout the nucleus, forming numerous secondary foci (**Fig. 1B**). A similar pattern of Bru distribution could be detected in myoblast-supplied myonuclei, albeit at a much smaller scale (**Fig. 1B**, also see **Fig 3**).

**Figure 2:**
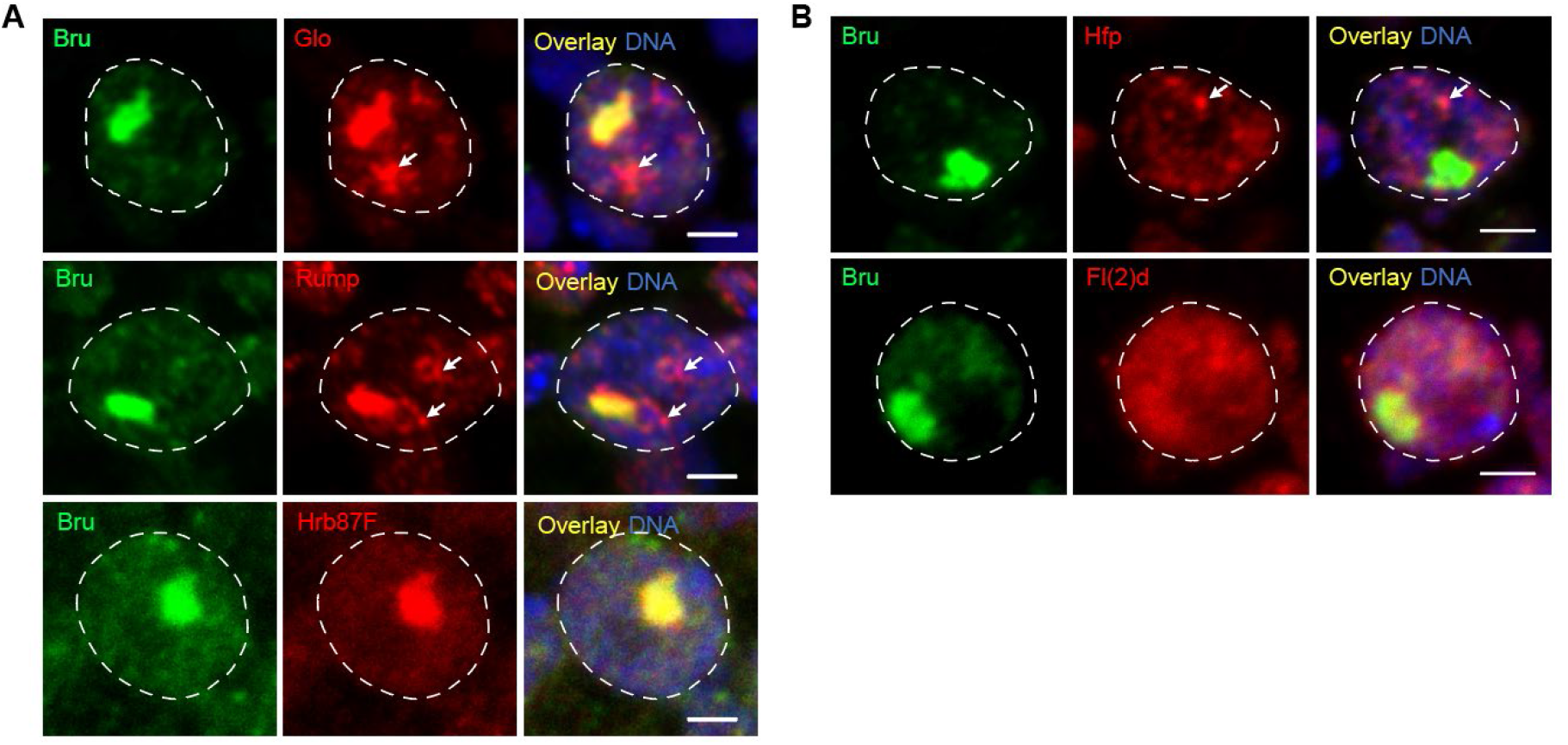
Protein composition of B-bodies. **A**. Proteins that co-localize with B-bodies. **B**. Proteins that do not co-localize with Bru or B-bodies. All images were obtained from 16-hour pupae using confocal microscopy, shown as maximum intensity projections of optical sections spanning entire nuclei. Co-localization is indicated by yellow signal in merged images. Arrows highlight regions of protein accumulation that do not co-localize with Bru. All proteins were detected by double immunofluorescence, except for GFP-tagged Hrb87F, which was expressed from a cloned genomic region and visualized via native fluorescence. Scale bar (all panels) = 2 µm.

**Figure 3:**
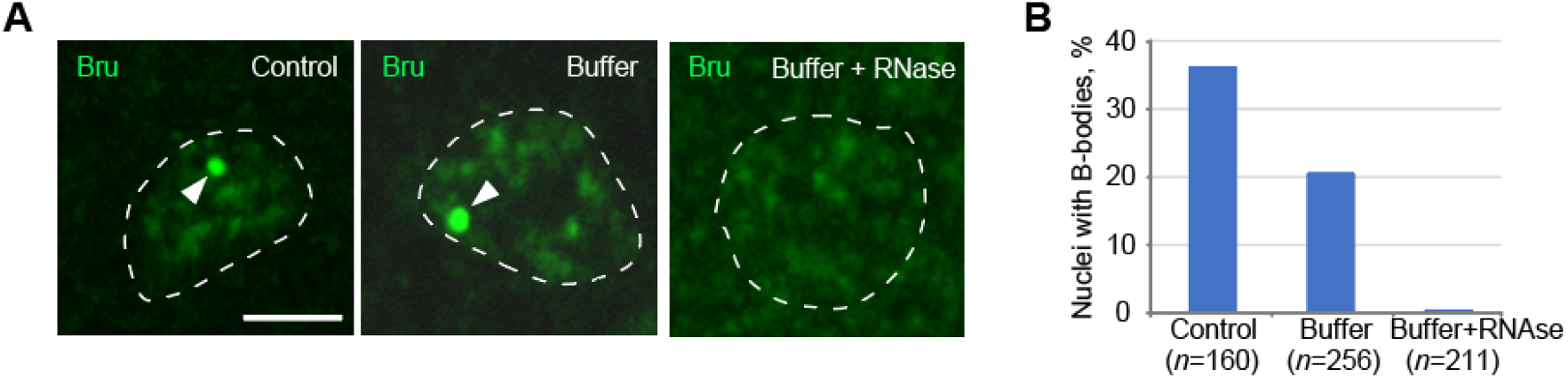
B-body integrity is sensitive to RNase treatment. **A**. Single-plane confocal images of DLM nuclei following different treatments. In the *Control* condition, tissue sections were fixed immediately. In the *Buffer* condition, samples were lysed with CSK buffer prior to fixation. In the *Buffer + RNase* condition, samples were lysed in CSK buffer supplemented with 1 µg/mL RNase A. Arrowheads indicate B-bodies; dashed lines outline nuclear boundaries. Scale bar = 2 µm. **B**. Quantification of B-bodies from the samples shown in panel A. *n* indicates the number of nuclei analyzed.

Another RBP that is known to accumulate in the B-body at 16 h apf is Muscleblind (Mbl) [3]. However, we observed that Mbl completely disappears from both B-body and DLM nuclei by 24 h apf **(Fig. 1C)**.

Recognizing that both Bru and Mbl are RBPs and splicing regulators, we have probed B-bodies for the presence of other proteins of similar functions. Double immunofluorescence revealed that Glo and Rump strongly accumulated in B-bodies where they co-localized with Bru (**Fig. 2A**). However, these two proteins were also found concentrating in nuclear foci outside B-bodies with partial or no co-localization with Bru. The splicing factor Hrb87F was tested via its GFP-tagged allele. The expression of Hsr87F was low, but the protein was present exclusively in B-bodies (**Fig. 2A**).

Some tested candidates were not B-body selective. For example, RBPs Hfp and Fl(2)d had moderate nuclear expression and demonstrated distinct patterns in which one protein (Fl(2)) was diffuse and the other (Hfp) formed nuclear speckles, but they did not concentrate in B-bodies (**Fig. 2B**). The expression of Orb2, the last tested RBP, was very weak and not conclusive (data not shown).

Our results significantly expand the list of B-body resident proteins. The identification of various RBPs in the B-body implies that this nuclear domain may contain an RNA component. Among all RPBs tested, Bru remains the strongest B-body marker with high expression levels and high fidelity to the B-body.

### B-body contains an RNA component

To further elucidate if the B-body has RNA in its composition, we treated unfixed the nuclei of flight muscles with RNase. We used serial cryosections of adult flies for this experiment to gain a better statistical power, since a single mature DLM contains hundreds of myonuclei as opposed to just a few myonuclei in LOMs. In control untreated sections, about 36.3% of analyzed nuclei (n=160) contained a distinguishable, although small, B-body (**Fig. 3**). A pre-treatment of unfixed cryosections spared most of B-bodies (occurrence 20.7%, n=256), while a combination of the lysis buffer with RNAse removed virtually all B-bodies from myonuclei (occurrence 0.5%, n=211) (**Fig. 3**).

These results strongly suggest that B-body contains RNA that may work as a structural component of this ND.

### Identifying hsrω in the B-body

To identify which specific RNA contributes to B-body organization, we performed a small-scale screening among lncRNA genes that produced transcripts longer than 5 kb and contained multiple clustered Bru-binding sites in the sequence (**Fig. 4A**, see Methods for selection details).

**Figure 4:**
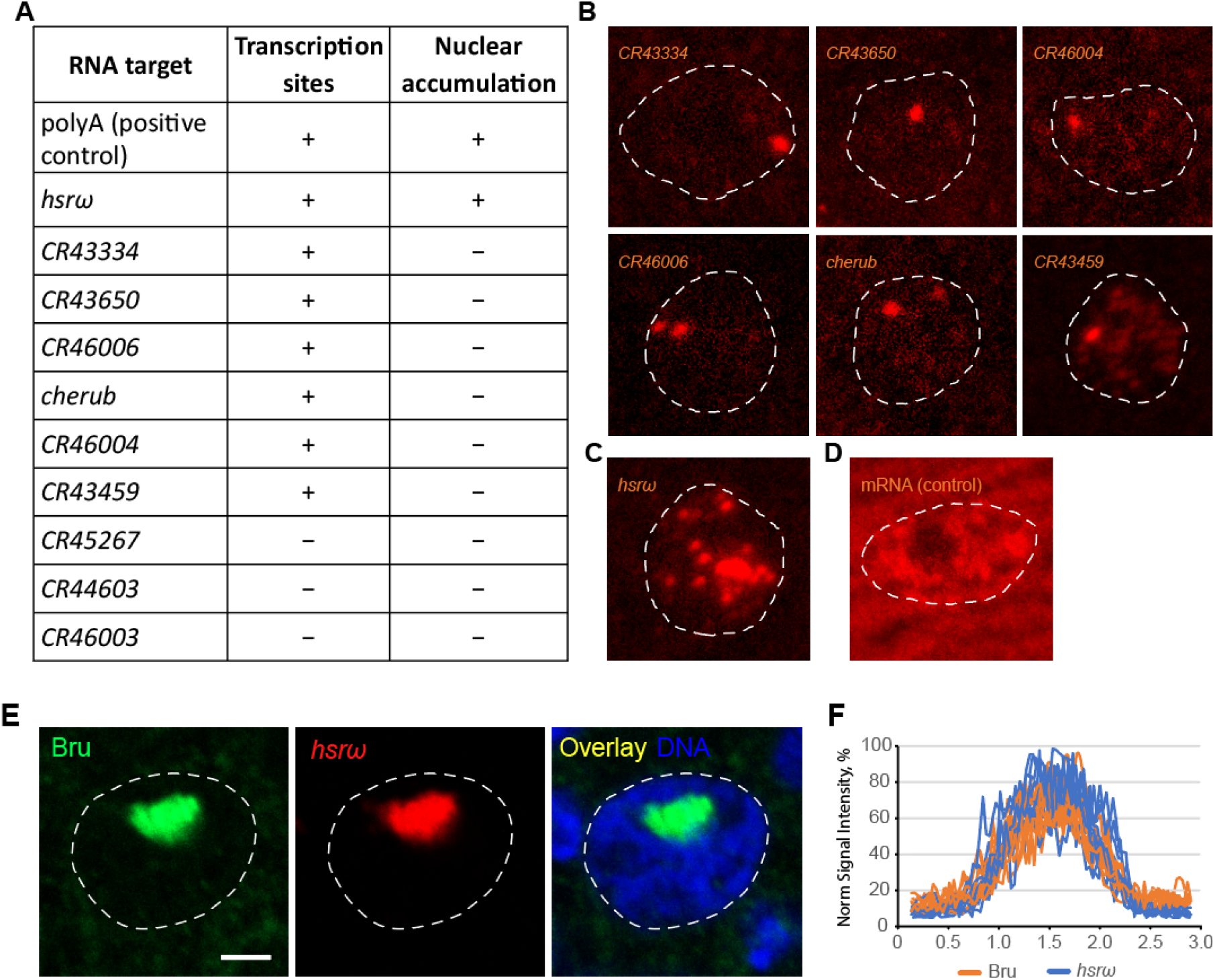
Identification of *hsrω* as a B-body RNA component. **A**. List of lncRNA genes included in the screen and corresponding outcomes. **B**. Examples of genes with positive expression but no nuclear accumulation. FISH signals are limited to transcription sites within the nucleus. ***C***. *hsrω* exhibits prominent nuclear accumulation resembling B-bodies, along with numerous smaller speckles. **D**. FISH positive control using a polyT probe, showing abundant signal in both nucleus and cytoplasm. **E**. Co-localization of *hsrω* (red) and Bru protein (green) at a B-body in a 16-hour pupa, revealed by combined IF-FISH. Scale bar = 2 µm. **F**. Intensity profiles of *hsrω* and Bru signals from 10 B-bodies, demonstrating complete co-localization. Intensity profiles were obtained by drawing a line across B-bodies in the Zen program.

The candidates were tested by FISH in developing DLMs at 48 h apf. At this time point, the density of sectioned muscles and the spacing between myonuclei were optimal to produce the best signal-to-noise ratio. Most of the tested lncRNAs displayed low nuclear expression, concentrating around their transcriptional sites (**Fig. 4B**); three candidates (*CR45267, CR44603*, and *CR46003*) were not expressed at all (**Fig. 4A**). One notable exception was the *hsrω* gene, which showed robust nuclear accumulation (**Fig. 4C**). The specificity of probe hybridization protocol was confirmed by control T30 probe that targeted the poly-A tails of mRNAs and produced ubiquitous signal from both nuclear and cytoplasmic compartments, as expected (**Fig. 4D**).

When the *hsrω* probe was applied to LOM nuclei at 16 h apf via ImmunoFISH, it strongly accumulated in B-bodies (**Fig. 5E**). *Hsrω* and Bru signals completely co-localized, confirming that *hsrω* is the constitutive component of the B-body (**Fig. 5F**).

**Figure 5:**
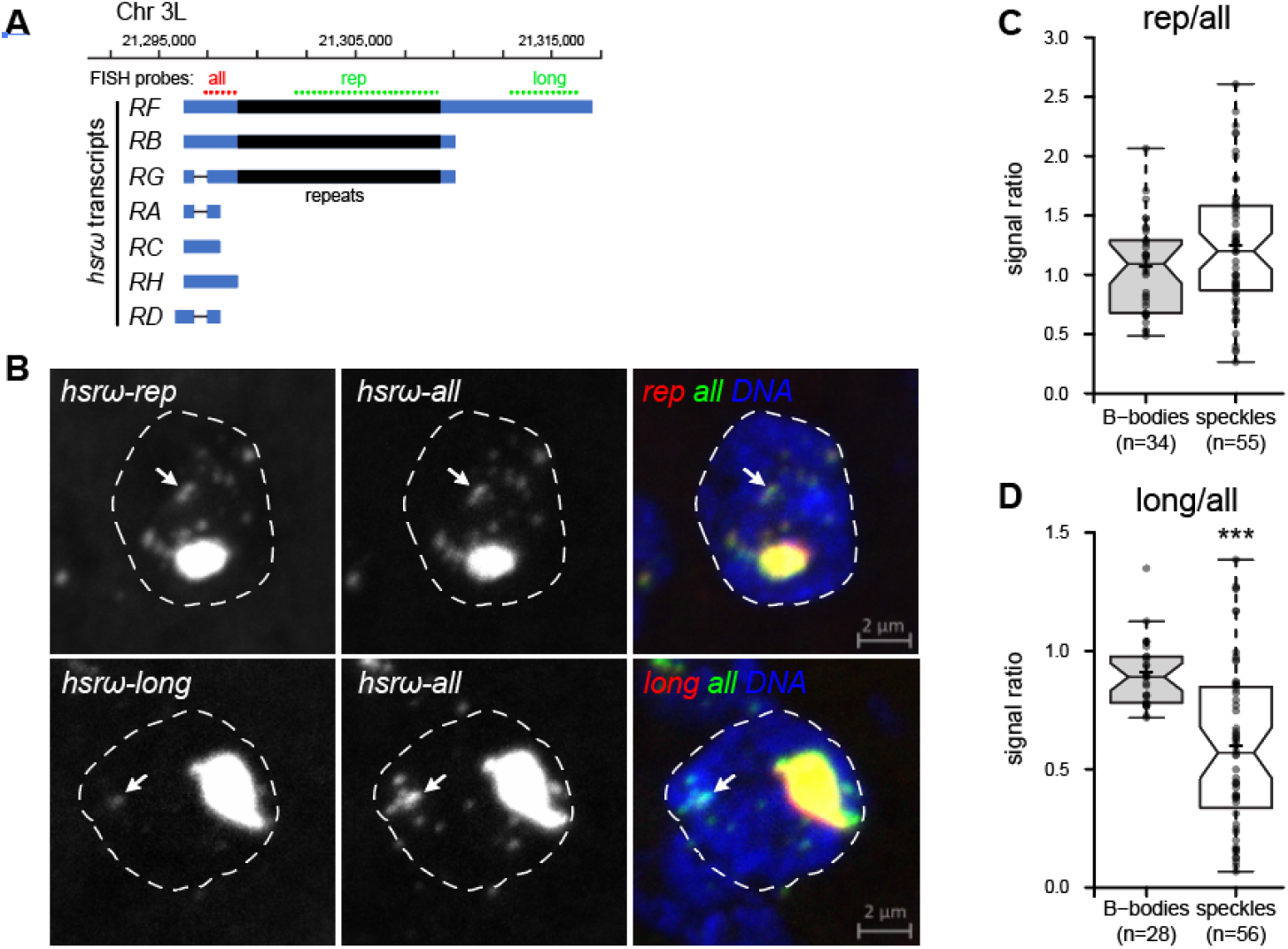
Isoform-specific localization of *hsrω* in B-bodies and omega speckles. **A**. Composition of the *hsrω* locus with FlyBase-annotated transcripts. Stellaris FISH probe target regions are indicated with dashed lines; the repetitive region is shown in black. **B**. FISH using different probes in LOM nuclei at 16 h apf. Grayscale individual channels are pseudocolored in merged images. Arrows indicate prominent omega speckles. The *hsrω-long* probe labels B-bodies well but shows weaker signal in omega speckles. **C, D**. Box plots of normalized FISH intensities for the *hsrω-rep* (C) and *hsrω-long* (D) probes, relative to *hsr-all* in B-bodies and speckles. Note reduced *hsrω-long* signal in speckles. Paired Student’s t-test; ***p < 0.001. *n* = number of analyzed structures.

### Isoform composition of B-bodies

The *hsrω* genetic locus produces multiple isoforms that vary in size and composition (**Fig. 5A**). The length of the short *hsrω* isoforms (annotated by FlyBase as *RA, RC, RH, RD*) is under 2.5 kb. Two medium-sized isoforms (*RB* and *RG*) have a length of ∼14 kb and feature a long region of tandem repeats [17]. The longest isoform *RF* is over 20 kb in length and, in addition to the repetitive region, features a 7.5 kb-long 3’ tail [17]. To better understand isoform composition of the B-body, we used double-color FISH with three different sets of probes targeting the common 5’ part of *hsrω* transcripts (hsr-all), the repeats (*hsr-rep*), and 3’ unique tail of the RF isoform (*hsr-long*). We observed that all probes have strong staining of B-bodies (**Fig. 5B**).

Nuclear distribution of *hsrω* in various tissues is often represented by small and numerous omega speckles [12]. In LOM myonuclei, multiple omega can be detected in addition to B-bodies, although their signal intensity by an order of magnitude was lower than that of B-bodies (**Fig. 5B**). *hsrω* transcripts containing the repetitive region were equally present in B-bodies and omega speckles, the hsr-rep probe was completely co-localized with hsr-all at (**Fig. 5B**,**C**). In contrast, the longest RF isoform was underrepresented in omega speckles (**Fig. 5 B,D**). The latter indicates that *hsrω* RF isoform may play a special role in the organization of B-bodies.

### Bru is dispensable for B-body structural integrity

In order to functionally test the role of protein component in the integrity of B-bodies, we applied RNAi to downregulate the canonical B-body resident, the Bru protein. The bru1 gene, expressing Bru, is not active in muscles besides IFMs, which allowed us to prevent Bru appearance in developing IFMs by running RNAi in LOMs using the Mef2-Gal4 driver. At 16 h apf, Bru could not be detected in LOM nuclei using immunofluorescence (**Fig. 6A**). However, the lack of Bru did not affect the appearance of B-bodies, that can be revealed by *hsrω* FISH (**Fig. 6A**). A quantification analysis of B-body area confirmed no changes in B-body sizes (**Fig. 6B**). Based on these results, Bru is a dispensible component of the B-body and does not have a structural significance for this ND.

**Figure 6:**
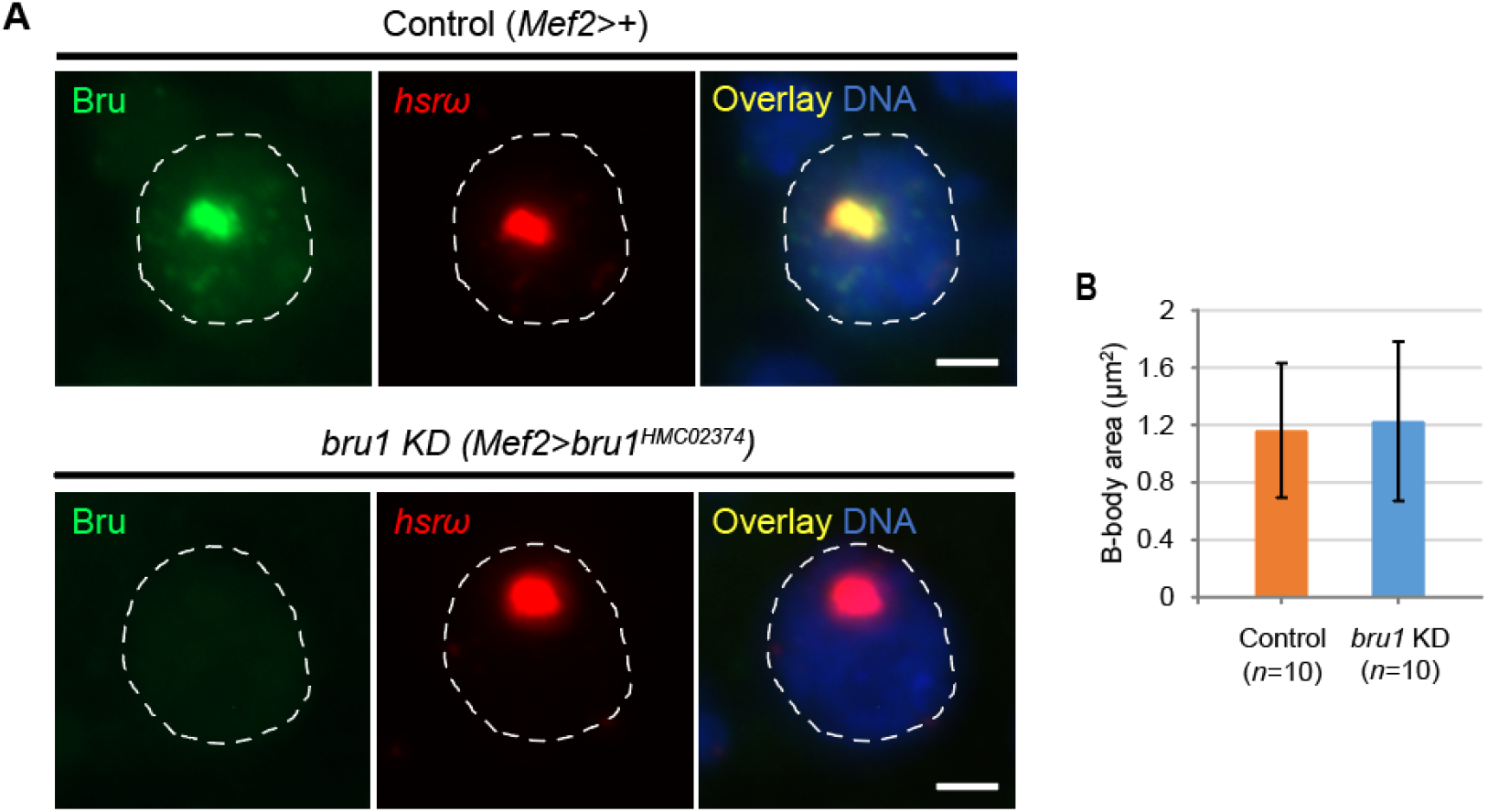
Bru is dispensable for B-body integrity. **A**. B-body appearance in 16 h apf pupae, revealed by double staining for Bru protein (green, immunofluorescence) and *hsrω* RNA (red, FISH). Bru knockdown (KD) was achieved by expressing RNAi against *bru1* under the muscle-specific *Mef2-Gal4* driver. Bru protein levels were reduced in KD samples, but B-bodies remained detectable via *hsrω* staining. Overlayed images show co-localization (yellow) in control nuclei. DNA is counterstained in blue. White dashed lines outline nuclear boundaries. Scale bars = 2 µm. **B**. Quantification of B-body size based on *hsrω* signal. Three biological replicates (pupae) were analyzed per condition; *n* indicates total number of LOM nuclei assessed. No significant difference was observed between control and KD samples (Student’s t-test, p > 0.05).

### hsrω is a structural scaffold of B-bodies

Next, we sought to determine the role of *hsrω* in the organization of B-body. Since RNAi was not effective in depleting the nuclear pool of *hsrω* in LOMs (data not shown), we employed a combination of genetic deficiencies to disrupt *hsrω*. The molecularly defined *hsr*-*ω*^66^ allele has a 1.6 kb deletion that removes the basal promoter, TSS, and extends into the first 8 bp of the *hsrω* gene [19]. This small deficiency was paired in *trans* with a larger deficiency *ED6052* that removes the *hsrω* gene but retains its upstream regulatory region intact (**Fig. 7A**). The Hsr66/ED6052 trans-heterozygotes developed normally, without any notable adverse phenotype. In addition, *ED6052* was combined with a larger chromosomal deficiency GC14 that has molecularly undefined breakpoints but phenotypically disables *hsrω* expression [20]. This combination of deficiencies was sublethal and adult Δ*hsrω*^GC/ED^ heterozygotes were rare.

**Figure 7:**
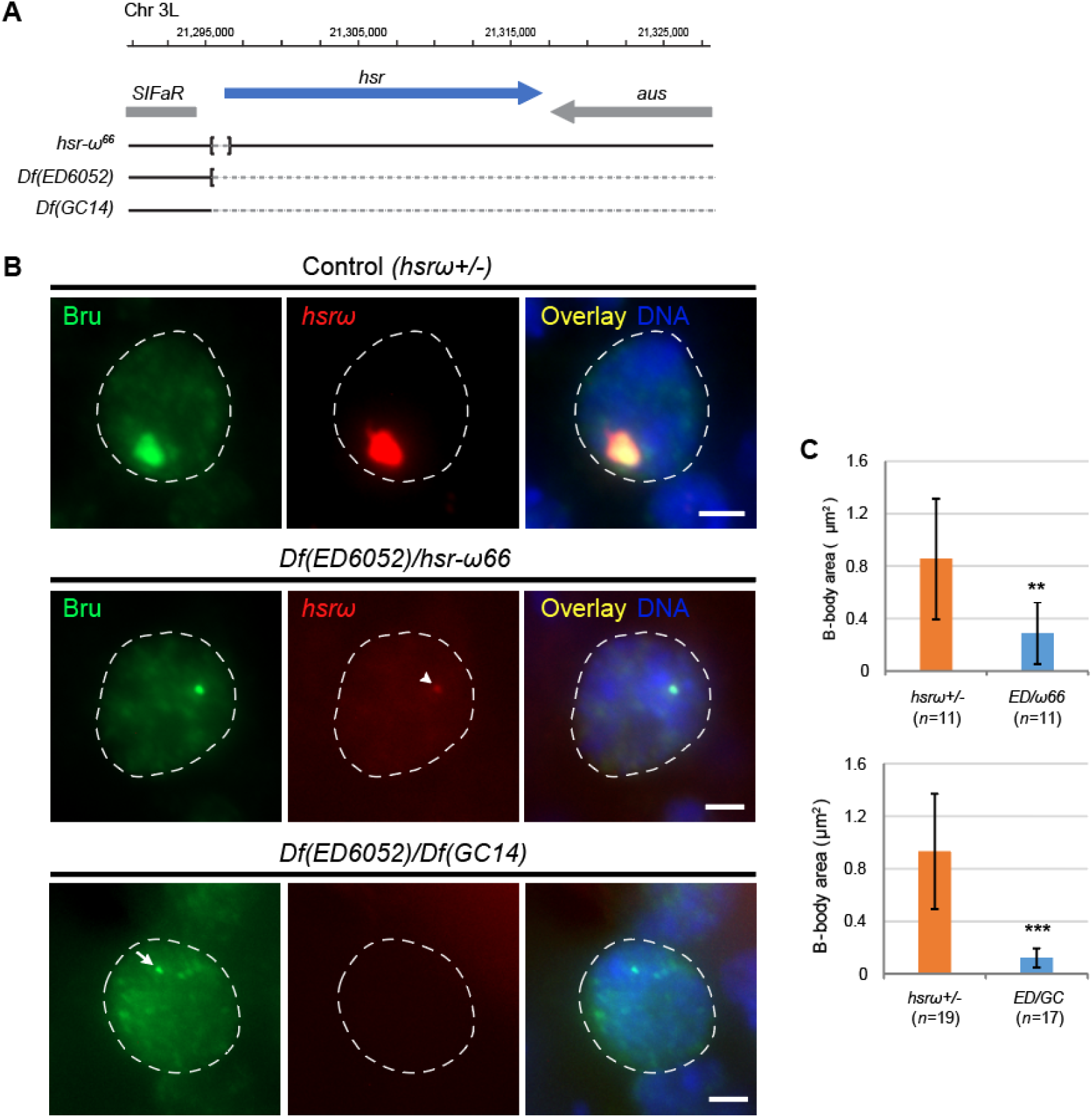
*hsrω* is an essential scaffolding component of the B-body. **A**. Schematic of genetic deficiencies used to delete the *hsrω* locus. Deleted regions are shown as dashed lines; square brackets indicate molecularly defined breakpoints. **B**. IF-FISH staining of LOM nuclei at 16 h apf. Controls are heterozygous pupae carrying one copy of a deficiency. Deficiency combinations caused a dramatic reduction of *hsrω* signal in the nucleus, though residual amounts were occasionally detected (arrowhead). In the absence of *hsrω*, Bru appears more diffuse, with occasional inclusions lacking *hsrω* signal (arrow). **C**. Quantification of B-body size based on Bru immunofluorescence. A significant size reduction was observed (Student’s unpaired t-test; **p < 0.01, ***p < 0.001).

B-body presence in pupae with *hsr* deficiencies was tested by FISH-IF. At 16 h apf, LOM nuclei of Δ*hsrω* ^ED/66^ flies had mainly diffuse Bru, with occasional small foci, which was in a striking contrast to the typical B-bodies that can be seen in heterozygous Δ*hsrω*^*+*/-^ siblings from the same cross (**Fig. 7B**). However, traces of *hsrω* expression could be still detected in some Δ*hsrω*^ED/66^ samples, and in such cases *hsrω* co-localized with small foci of Bru (**Fig. 7B, arrowheads**). Alternative trans-heterozygotes Δ*hsrω*^GC/ED^ had no detectable *hsrω* FISH signal. In these pupae Bru nuclear diffused distribution, with occasional speckles (**Fig. 7B**). We used Bru immunofluorescence to quantify residual B-bodies or B-body-like structures. By quantitative analysis, both combinations of genetic deficiencies produced significantly smaller Bru foci, with the *ED6052*/*GC14* having the most significant impact (*p*<0.01) **(Fig. 7C)**.

These results demonstrate that lncRNA *hsrω* is essential for the structural organization of B-body and can be considered a scaffolding compent of this nuclear domain. However, elimination of B-bodies did not eliminate Bru as it was easily detectable in the nuclei. ability of Bru to form foci, albeit severely reduced, without *hsrω* suggests some degree of protein contribution in B-body making.

### Bru is a nonessential but impactful component of the B-body

In *hsrω* depleted nuclei, Bru was often found in small but compact inclusions that appeared as protein aggregates (**Fig. 7B, arrow**). This may suggests that *hsrω* functions in B-bodies as a stabilizing factor that prevents aggregation at high local concentrations of proteins. To experimentally test this hypothesis, we altered the protein-RNA balance in the nucleus by ectopically overexpressing Bru. To overcome early lethality associated with Bru overexpression, we used the temperature-sensitive Mef2TS-Gal4 driver, which was activated in 3^rd^ instar larvae, shortly before pupation. A GFP-tagged Bru was traced to the LOM nuclei at 16 h apf, where it accumulated thoughout the nucleus and often was found in compact inclusions that did not colocalized with *hsrω* (**Fig. 8A**). The frequency of nuclear aggregates in Bru-overexpressing nuclei was high and in a comparable range with the frequencies of Bru aggregates in hsr-depleted nuclei (**Fig. 8B**). This finding provides an important insight into the functioning of the B-body and a depot for RBPs.

**Figure 8:**
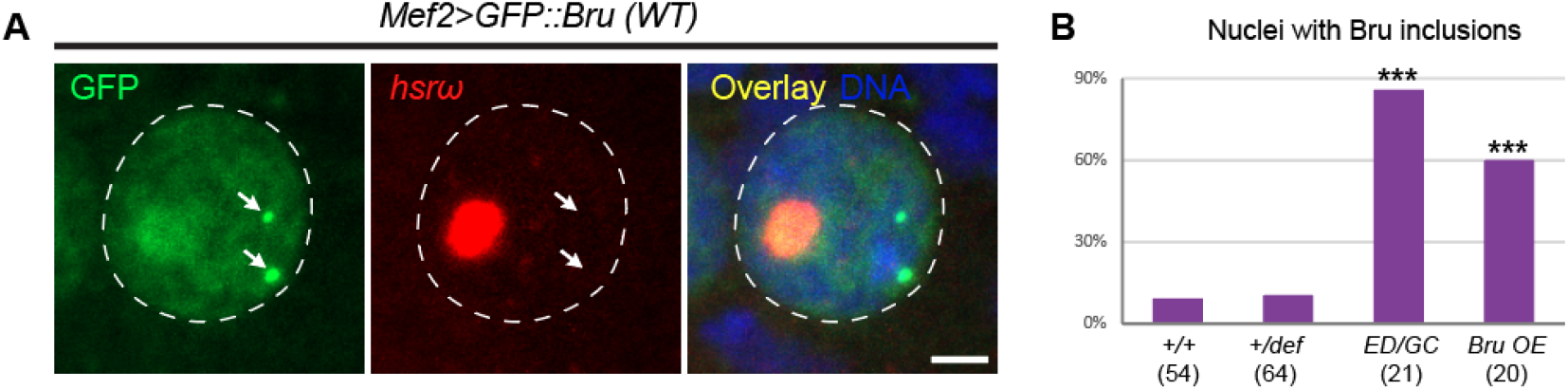
Imbalance between Bru and *hsrω* leads to protein aggregation. **A**. IF-FISH staining of LOM nuclei at 16 h apf overexpressing GFP-tagged Bru. Arrows mark nuclear inclusions lacking *hsrω* signal. **B**. Quantification of Bru-positive inclusions in control, *hsrω*-depleted (ED/GC), and Bru-overexpressing nuclei. Deficiency labels follow those used in Fig. 7. *n* indicates the number of nuclei analyzed per group. Statistical analysis by Fisher’s exact test; ***p < 0.001.

## DISCUSSION

In this study we have analyzed the composition of the B-body, a novel and previously uncharacterized ND. We demonstrate that the B-body accumulates multiple RBPs, and we identified lncRNA *hsrω* as a critical structural component. Our findings imply that B-body protects sequestered proteins by preventing their aggregation, and *hsrω* plays a pivotal role in this process.

### B-bodies and omega-speckles

*Hsrω* has been in the focus of research for a long time, but most of the attention was drawn to small and numerous nuclear domains known as omega speckles [12]. The larger site of *hsrω* concentration was regarded merely as *hsrω* transcription site, which corresponded to the gene’s position at the cytoband 93D4 on 3R polytene chromosome [16]. The B-bodies that we observe in the developing IFMs is, technically, a greatly expanded *hsrω* transcription site, with the size on par with the biggest nuclear domain, the nucleolus [4]. This enormous size of the B-body that dwarfs in comparison transcription sites of actively transcribed genes suggests the presence of a specific consolidating factor that immobilizes *hsrω* transcripts and prevents their diffusion upon transcription. Our data implies that such factor could be the longest RF isoform. This isoform runs >20 kb and includes the unique, ∼5 kb tail that is not present in other isoforms (**Fig. 5A**). According to our FISH data, the RF tail is enriched at B-bodies but depleted in omega speckles. Although direct measurements could not be made, the irregular size and quantities of omega speckles argues in favor of the high dynamic nature of these structures, as compared to the stable and less diverse morphology of B-bodies. Since the occurrence of the repetitive region does not diminish in omega speckles, a likely candidate is the long and unique tail of the RF isoform. In line with these findings is that under stressful conditions the expression of RF isoform steeply increases and that coincides with expansion of transcription site on polytene chromosome and reduction (to a complete elimination) of the omega speckles [16]. Whether the RF tail directly stabilized *hsrω* transcripts by RNA-RNA or steric interactions, or it does so via protein components remains to be determined.

Considering the stress-activated expression of the RF isoform, it is tempting to speculate that B-bodies are initially formed in remodeled LOMs (see Fig.1) in response to the stress caused by atrophic changes that take place in these muscles during early metamorphosis. Interestingly, transcription sites of *hsrω* in IFM nuclei acquired from myoblasts remain of normal and not enlarged.

### B-body analogues in other organisms

In general, lncRNAs have poor sequence conservation across even closely related species complicates identification of homologous genes by sequence comparison, but there are several striking homologs that share common on characteristic features, namely long size, interaction with RNA proteins, and involvement in nuclear domains. Two such functional homologs are MALAT1 in mammals and Satellite III (SatIII) transcripts in humans.

MALAT1 is a lncRNA of approximately 8 kb that is abundantly expressed and localized in the nuclei of mammalian cells. It forms distinct foci that colocalize with nuclear speckles, domains rich in splicing factors like SR proteins [21]. This suggested an architectural role for MALAT1 like that of *hsrω* in forming omega speckles and B-bodies in Drosophila. However, the structural significance of MALAT1 remains somewhat ambiguous. Its recruitment to nuclear speckles is delayed after mitosis, indicating that speckle formation precedes MALAT1 accumulation [21].

A closer functional homolog to *hsrω* in humans is the SatIII RNA, which is induced by heat shock, much like *hsrω* [22]. SatIII transcripts are produced from pericentromeric repetitive sequences and accumulate at their transcription sites to form nuclear stress bodies NSB. These temporary nuclear domains recruit and sequester RBPs, including splicing regulators such as SF2/ASF and SRp30c [22]. The presence of repeats in SatIII RNA, accumulation of RBPs, and temporary nature of NSB runs a close parallel with *hsrω* and B-bodies. To continue this analogy, we may assume that B-bodies are formed as part of the stress response during atrophic processes that take place in LOMs as they undergo atrophy before transitioning into IFMs [23].

### The dynamic protein content of the B-body

In this study we demonstrate that in addition to Bru, the B-body is populated by multiple proteins. Interestingly, we found evidence that B-body residents demonstrate mobility by leaving this nuclear domain, which was the case for Mbl (**Fig. 1**). These dynamics argue in favor of the protein exchange between this nuclear domain and the cytoplasm, instead of being permanently trapped. We hypothesize that Bru replaces Mbl after competing for the same mRNA binding sites to provide flight-muscle-specific mRNA splicing, making IFMs different from other muscles where Mbl is continuously expressed [24]. This notion is opposite to the situation reported for pathological RNA inclusions human patients of myotonic dystrophy [25]. Under these conditions human Mbl homologue MBNL1 becomes trapped in these inclusions permanently. Understanding how B-body residents retain mobility might help resolve the pathology of myotonic dystrophy.

As flight muscle develops, Bru becomes less concentrated within B-bodies as it spreads throughout the nucleus starting from 24 h apf (**Fig. 1**). The mechanism of such a transition is currently unknown and requires more research. A previous study showed a dramatic effect of phosphorylation on the integrity of another type of nuclear domain – nuclear speckles [26]. Similarly, phosphorylation by protein kinases may be required to regulate the affinity of Bru to the B-body, and was shown to be phosphorylated *in vitro* [27]. Phosphorylated Bru might lose affinity to *hsrω* and dissociate from the B-body.

## Notes

### Competing Interest Statement

The authors have declared no competing interest.

### Funding

- National Science Foundation: 2042814

